# Soybean Haplotype Map (GmHapMap): A Universal Resource for Soybean Translational and Functional Genomics

**DOI:** 10.1101/534578

**Authors:** Davoud Torkamaneh, Jérôme Laroche, Babu Valliyodan, Louise O’Donoughue, Elroy Cober, Istvan Rajcan, Ricardo Vilela Abdelnoor, Avinash Sreedasyam, Jeremy Schmutz, Henry T. Nguyen, François Belzile

## Abstract

Here we describe the first worldwide haplotype map for soybean (GmHapMap) constructed using whole-genome sequence data for 1,007 *Glycine max* accessions and yielding 15 million variants. The number of unique haplotypes plateaued within this collection (4.3 million tag SNPs) suggesting extensive coverage of diversity within the cultivated germplasm. We imputed GmHapMap variants onto 21,618 previously genotyped (50K array/210K GBS) accessions with up to 96% success for common alleles. A GWAS performed with imputed data enabled us to identify a causal SNP residing in the *NPC1* gene and to demonstrate its role in controlling seed oil content. We identified 405,101 haplotypes for the 55,589 genes and show that such haplotypes can help define alleles. Finally, we predicted 18,031 putative loss-of-function (LOF) mutations in 10,662 genes and illustrate how such a resource can be used to explore gene function. The GmHapMap provides a unique worldwide resource for soybean genomics and breeding.

## Introduction

Soybean (*Glycine max* [L.] Merr.) is a unique crop with substantial economic value. It is the largest plant source of both animal feed protein and edible oil. It also plays a key role in sustainable agriculture as it fixes atmospheric nitrogen with the help of microorganisms (Hymowitz 1970). Diverse evolutionary processes and forces (including cycles of polyploidization and subsequent diploidization), along with domestication and modern breeding have shaped the soybean genome (Schmutz et al. 2010). The detection of the molecular footprints of these processes is essential for understanding how genetic diversity is generated and maintained and for identifying allelic variants responsible for phenotypic variation (Torkamaneh et al. 2018).

The global production of soybean has increased substantially in recent years (**Supplementary Figure 1**), but the rate of annual yield gains has lagged behind that of maize (FAOSTAT Database). In addition, with increased fluctuations in climatic conditions, next-generation soybean cultivars must not only be higher yielding but also more resilient to multiple abiotic and biotic stresses (Djanaguiraman et al. 2018). In the main soybean-growing areas of the world, soybean is an introduced crop and the foundational germplasm was very limited in its genetic diversity (Hyten et al. 2006; Maldonado dos Santos 2016). Continued genetic improvement in soybeans will require a better understanding of the genetic and especially allelic diversity within worldwide resources (Qiu et al. 2013).

Here we present the first haplotype map for soybean (GmHapMap) assembled from DNA resequencing data for a collection of 1,007 worldwide *G. max* accessions. We explore the use of this GmHapMap for (i) imputation of untyped variants to create high density genotype data required for gene-level resolution of genomewide association studies (GWAS); (ii) construction of gene-centric haplotypes (GCHs) for the entire set of soybean genes; and (iii) identification of 11K knock-out genes due to loss-of-function (LOF) mutations. The GmHapMap provides a unique resource for translational and functional genomics for the worldwide soybean community.

## Results

### Development of GmHapMap

#### Genomic variation

To establish a first worldwide haplotype map for soybean (GmHapMap), a total of 1,007 resequenced soybean accessions, representative of the worldwide cultivated germplasm, were used (**Figure 1A**). These accessions span thirteen maturity groups (MGs) (000-X) based on their latitudinal adaptation. This collection includes 727 previously resequenced accessions, as well as 280 accessions sequenced as part of this study which were selected to achieve a more complete coverage of soybean worldwide diversity. Genome sequencing, analyses, and accession information for GmHapMap accessions are summarized in a **Supplementary Note** and **Supplementary data 1**.

In total, 165 billion paired-end reads (100-150 bp; total of 19.2 trillion bp) provided an average depth of coverage of more than 15× and these were analyzed using a single pipeline (Fast-WGS) to ensure uniform variant calling. After mapping against the soybean reference genome (cv. Williams 82 va2.v1) (Schmutz et al. 2010), we identified 14,872,592 nucleotide variants (**Table 1**), including 13M single- and multiple-nucleotide variants (SNVs and MNVs) and 2M small insertions/deletions (InDels) (−50 bp to +32 bp). Approximately 45% of these were rare (minor allele frequency (MAF) < 5%) (**Supplementary Figure 2**). Coding regions represent ~6% of the soybean genome, but only ~2.3% of the total nucleotide variants were present in these regions (**Table 1**) with an average non-synonymous/synonymous ratio of 1.49. Nucleotide variants were 2-fold more abundant in coding regions compared to InDels, however InDels were overrepresented in the regulatory regions. Missing data comprised less than 8% of the data, and these were subsequently imputed with high accuracy (*r^2^* = 99.7%). Using independent genotyping data (SNP array and dbSNP database), we estimated the false-positive rates of nucleotide variants to be ~0.03%. This constitutes an extensive and highly accurate set of foundational data for a soybean haplotype map.

**Table 1.** Type, number and location of nucleotide variants in GmHapMap.

| Statistics |  | Location (%) |  |  |  |  |  |
| --- | --- | --- | --- | --- | --- | --- | --- |
| Types | Count | CDS* | Intron | UTR† | Splice sites | Up/Down stream‡ | Intergenic |
| SNV | 12,197,920 | 2.5 | 8.5 | 1.6 | 0.2 | 44.8 | 42.4 |
| MNV | 801,373 | 2.3 | 6.9 | 1.0 | 0.1 | 44.6 | 45.1 |
| INS | 887,485 | 0.9 | 10.2 | 2.7 | 0.2 | 54.3 | 31.6 |
| DEL | 985,814 | 1.0 | 10.1 | 2.3 | 0.3 | 53.0 | 33.2 |
| **Total** | **14,872,592** | **2.8** | **8.6** | **1.7** | **0.2** | **45.9** | **41.7** |

\*Coding DNA sequence; †Untranslated region; ‡ 5 kb before or after a gene

#### Extensiveness of GmHapMap

The extensiveness of the GmHapMap was measured based on nucleotide diversity and haplotype diversity. Previously, the SoySNP50K array has been used to genotype the entire USDA soybean germplasm collection (20,087 accessions of *G. max* and *G. soja*) (Song et al. 2013). We found that GmHapMap includes nearly all polymorphisms (99.4%) with a MAF > 1%, as well as ~89% of rare SNPs (MAF < 1%) documented within these *G. max* accessions. Haplotype diversity (pairwise LD using both r^2^ and D’) was calculated for sequence variants and the average distance over which LD decayed to 0.2 was ~138 kb (**Supplementary Figure 3**). We identified 4.3 million haplotype-based tag SNPs and, to determine if a good level of saturation of both variants and haplotypes had been achieved, we randomly selected subsets of samples of increasing size (N=100, 200, …, and 1,007). As illustrated in **Figure 1B**, the number of variants discovered did not increase significantly beyond ~750 accessions, while the number of haplotypes reached a plateau much faster (within the first ~500-600 accessions). Together, these results suggest that the GmHapMap dataset offers an exhaustive characterization of the variants and haplotypes present in soybean germplasm.

#### Genetic diversity and artificial selection

Bayesian clustering (STRUCTURE) of GmHapMap accessions using whole-genome SNP data revealed 12 subpopulations (**Supplementary Note & Supplementary Figure 4**). We explored the phylogenetic relationships among GmHapMap accessions by constructing an un-rooted neighbor-joining tree. As can be seen in **Figure 1C**, the grouping of accessions reflected geographic origin with some admixture (**Figure 1C and Supplementary Figure 5**). Genomewide genetic diversity (θπ) analysis showed a consistent level of genetic diversity (mean of θπ = 1.36 × 10^−3^, ranging between 1.19 × 10^−3^ to 1.72 × 10^−3^) in different soybean populations (**Figure 1A**). Nucleotide diversity was plotted for the 20 chromosomes and found to be highest in the terminal regions of chromosomes (**Supplementary Figure 6**). Extensive peri-centromeric regions were very low in genetic diversity but chromosomes Chr13 and Chr17 maintained higher diversity across these regions. Genic regions showed even lower levels of diversity (mean θ_π_ = 7.1 × 10^−4^) and exceptionally low diversity was seen within 527 genomic regions (including 540 genes; mean θ_π_ = 4.6 × 10^−6^) that are presumably selection hotspots (**Supplementary Note, Supplementary data 2 and Supplementary Figure 7**).

**Figure 1.**
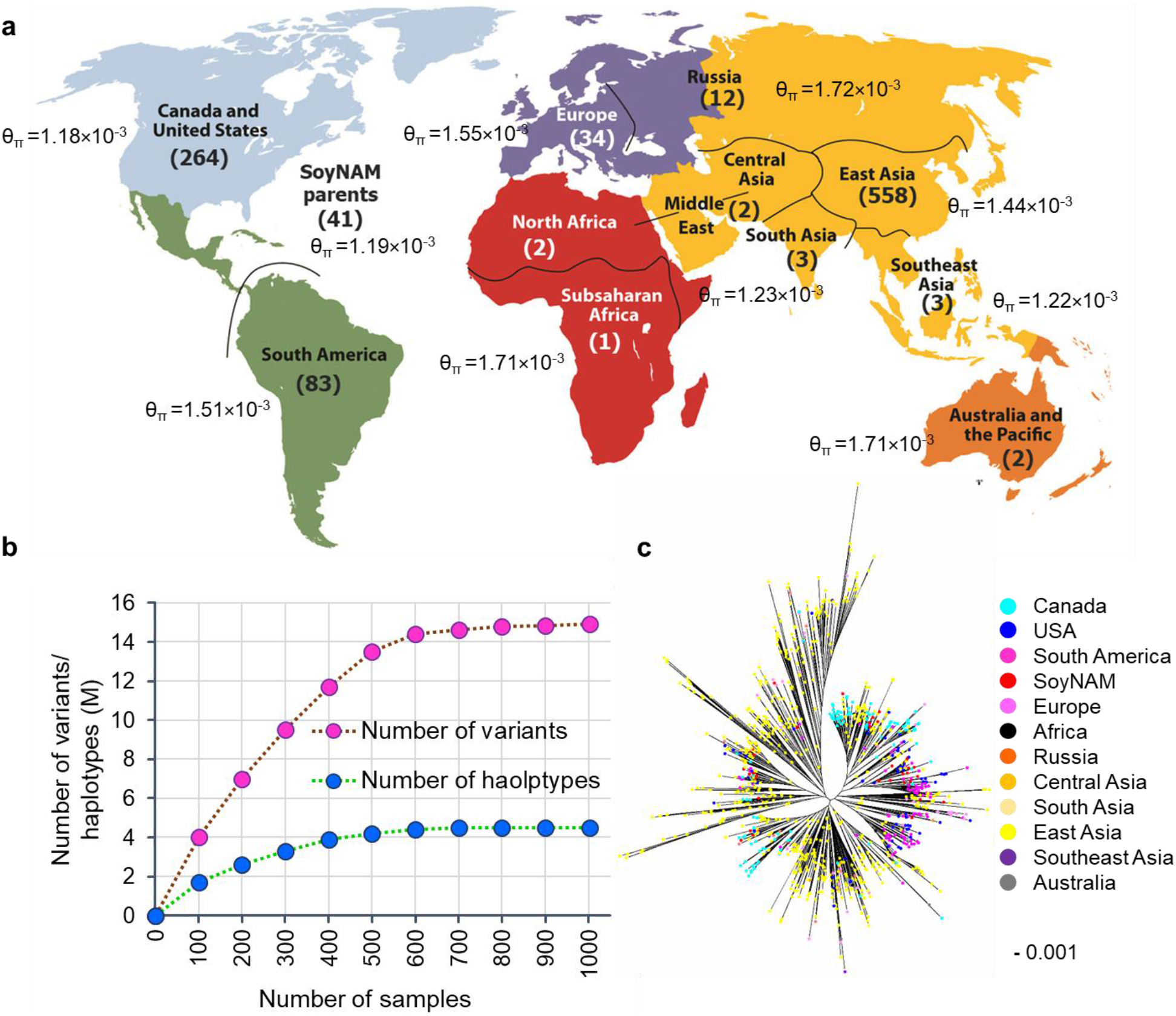
Description of GmHapMap. (a) Geographical distribution and related genetic diversity value (θπ) of GmHapMap accessions. (b) Number of variants (pink) and haplotypes (blue) based on different number of accessions. (c) Un-rooted phylogenetic tree of all accessions inferred from whole-genome SNPs representing existing genetic diversity and admixture among GmHapMap accessions.

### Phasing, identity-by-descent, and large-scale imputation of untyped variants

The GmHapMap dataset captures substantial amounts of identity-by-descent (IBD) allele sharing which allows a rule-based approach to long-range phasing that yields very accurate haplotypes. Using long-range phasing, we found 95 blocks of IBD larger than 1 Mb in size (**Supplementary data 3**). The determination of haplotype phase is important because of its applications such as the imputation of untyped variants. Imputation of untyped variants greatly boosts variant density, allowing fine-mapping studies of GWAS loci and large-scale meta-analysis. We created two reference panels: REF-I comprising all SNPs and REF-II containing 1.9M haplotype-based tag SNPs that reside in genic regions. Three lower density genotype datasets, SoySNP50K (20,087 accessions genotyped with 43K SNPs), genotyping-by-sequencing (GBS; 1,531 accessions genotyped with 210K SNPs), and combined GBS/SoySNP50K (1,531 accessions genotyped with 250K SNPs) were used for untyped variant imputation with each of the two reference panels. In all but one case, the accuracy (squared correlation (R^2^) between imputed and known genotypes, see M&M for details) ranged between 92% and 96% for common variants (allele frequency (AF) > 0.2) in each dataset, while decreasing gradually with allele frequency (**Figure 2A**). In the case of the SoySNP50K dataset using REF-I, the accuracy of imputed untyped variants was significantly lower (80-85% for common alleles). Given the observed variation in the accuracy of imputation using different reference panels and datasets, we investigated the causes of erroneous inferred calls. Several characteristics were tied to inaccurately imputed SNPs: these were commonly rare variants (low AF), located in recombination hot spots, in short LD blocks or in genomic regions with structural variants. Furthermore, the initial marker density in the experimentally-derived dataset had a large impact on imputation accuracy. GBS and SNP array datasets are two highly complementary marker datasets because most (~90%) of the SoySNP50K markers are present in genic regions, while most of the GBS markers (~60%) are present in intergenic regions (**Supplementary Figure 8 & 9**). Therefore, combining GBS and SoySNP50K datasets (**Supplementary Note**) increases the density and uniformity of distribution of SNPs across the genome. The joint use of such commonly available SNP data increased the level of accuracy of imputation of untyped variants (**Figure 2A**).

To demonstrate the benefits of untyped-variant imputation on GWAS analysis, the imputation was performed on a 1Mb-region harbouring a QTL previously identified for seed oil content on chromosome 14. We used the REF-II panel to perform imputation on an initial dataset of 64K GBS-derived SNPs (genomewide) among 139 soybean lines that had been characterized for their seed oil content (Sonah et al. 2015). Using this enhanced SNP catalog and a multi-locus mixed-model implementation, a very strong association (*p*-value = 4.2×10^−14^ and *q*-value < 0.001) with a SNP residing in the *NPC1* (Niemann-Pick C1) gene (*Glyma.14g001500*) (**Figure 2B**) was detected. An Arabidopsis mutant of this gene (*npc1*) exhibits a 58% higher fatty acid content (Feldman et al. 2015) making this gene a likely candidate contributing to total oil content in soybean. This demonstrates that the increased number of informative SNP loci, obtained through the imputation of untyped variants, can prove highly beneficial in studying the genetic architecture of complex agronomic traits in soybean.

**Figure 2.**
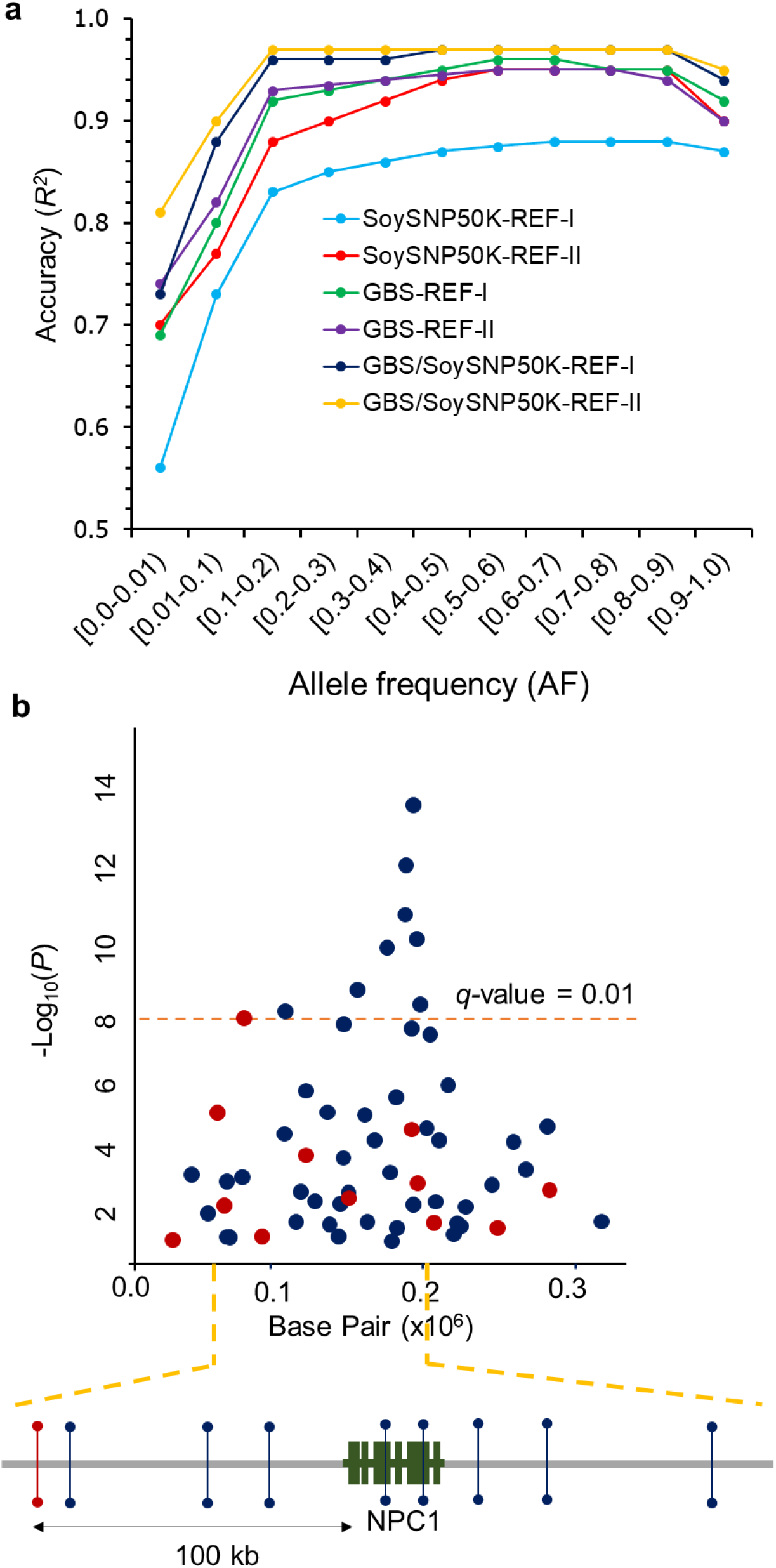
(**a**) Imputation accuracy as a function of allele frequency for 6 different scenarios; three different experimentally derived genotype datasets (SoySNP50K, GBS, and GBS/SoySNP50K) and two reference panels (REF-I and REF-II). (**b**) Top, association analysis for seed oil content on chromosome 14. Blue dots represent imputed variants whereas red dots identify the original GBS-derived variants. *NPC1* (Niemann-Pick C1) is an orthologue of an Arabidopsis gene known to play a key role in fatty acid synthesis. Bottom, schematic representation of strongly associated variants in the vicinity of the *NPC1*. The nearest significantly associated GBS-derived variant (red line) is located 100 kb upstream and exhibits a relatively low degree of LD (r^2^ = 0.5).

### Gene-centric haplotypes: a resource for translational genomics

HaplotypeMiner (Tardivel et al. unpublished) and the GmHapMap SNP dataset were used to identify 405,101 gene-centric haplotypes (GCHs) for 52,823 genes (94.5% of all soybean genes (55,589)). As can be seen in **Figure 3A**, the number of GCHs per gene ranged between 2 and 43, while averaging ~7 (**Supplementary Figure 10, and Supplementary data 4**). GCHs could not be determined (ND) for 2,766 genes with the set of parameters used here. In total, 11,407 genes had more than 10 GCHs with 71% (8,082 genes) of these harboring 11-15 GCHs. Such genes were typically located in very short LD blocks with a high degree of nucleotide diversity (mean θ_π_ = 4.5 × 10^−3^) (**Supplementary Figure 11**). A slight negative correlation was observed between gene length and the number of GCHs. However, we found a positive correlation between GCH counts and haplotype size (distance between two most distant SNPs defining a GCH) (**Supplementary Figure 12**). An example of GCHs for the *GmGIa* (*Glyma.10g221500*) gene (*E2* locus controlling maturity) (Watanabe et al. 2012; Tsubokura et al. 2014), an orthologue of the arabidopsis *GIGANTEA* (*GI*) gene, is presented in **Figure 3B**. We found three GCHs for *GmGIa*, which is consistent with the number of alleles that have been previously reported for this gene. Knowledge of the GCHs (and possibly alleles) in all soybean genes can greatly facilitate the establishment of a functional link between the various alleles of a gene and the associated phenotype.

**Figure 3.**
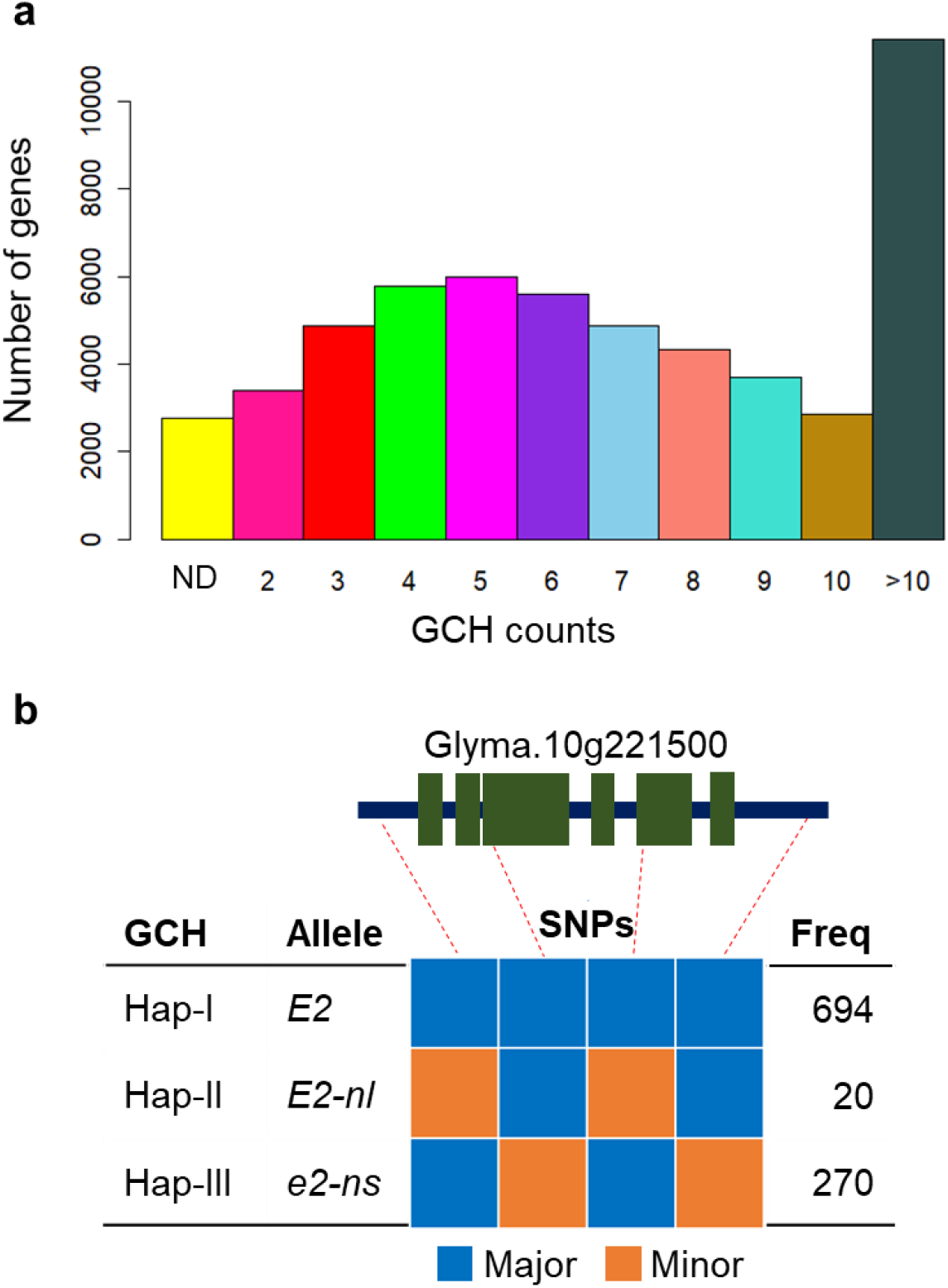
Description of GCHs characterized in GmHapMap dataset. (a) Distribution of number of genes based on their predicted GCHs. (b) Schematic representation of predicted GCHs for *GmGIa*.

### LOF Mutations: a resource for functional genomics

Using SnpEff, a subset of variants located inside the coding regions were predicted to have a large functional impact. Of these variants, 18,031 putative loss-of-function (LOF) mutations are predicted to severely impair protein synthesis or function through disruption of splicing, introduction of a premature stop codon, shifts in the coding frame and alterations to the start/stop codons (MacArthur et al. 2012) and these were identified in a total of 10,662 genes (19.3% of all soybean genes) (**Table 2**). These mutations are the result of 5,987 SNVs (33.2%), 279 MNVs (1.5%) and 11,765 InDels (65.3%). Frameshift-inducing variants (10,754) were the predominant category, representing 59.6% of LOF mutations and affecting 6,718 genes. InDels (ranging from −50 bp to +32 bp) were, understandably, over-represented (4-fold) in the LOF category due to their high probability of resulting in a LOF allele. Overall, most of the LOF mutations were present at low frequency, with 78% having an allele frequency below 10% (**Supplementary Figure 13**). Genes harboring LOF one or more mutations were categorized into two groups: unique and multi-copy. We reasoned that a LOF mutation in a unique gene would necessarily result in phenotypic consequences. We found that only 706 (6.6%) of genes were single-copy genes, while the remaining 9,957 (93.4%) had at least one other copy. This constitutes a significant enrichment (*P* < 0.001) compared to the genomewide occurrence of gene duplication. LOF mutations in duplicated genes could also have functional consequences if the mutated copy was uniquely expressed as a consequence of neo- or sub-functionalization (Roulin et al. 2013). We assessed this by examining transcriptomic data from 26 tissues and found that 9,570 of the 9,957 duplicated genes (96%) exhibited a unique expression pattern (**Supplementary Note, Supplementary data 5 & Supplementary Figure 14**). Thus, despite the fact that the vast majority of LOF mutations occur in genes for which there is more than one copy, a large proportion of these genes exhibit unique expression patterns, thus making it possible that a LOF will result in a detectable phenotype.

**Table 2.** Number of loss-of-function variants by sequence ontology (SO).

| SO term | SNV | MNV | INS | DEL | Total variants | Genes |
| --- | --- | --- | --- | --- | --- | --- |
| Splice site-disrupting (donor) | 1,270 | 38 | 247 | 205 | 1,760 | 1,640 |
| Splice site-disrupting (acceptor) | 1,546 | 52 | 207 | 146 | 1,951 | 1,803 |
| Stop codon-introducing | 2,826 | 149 | 100 | 7 | 3,082 | 2,418 |
| Frameshift-inducing | 0 | 0 | 4,158 | 6,596 | 10,754 | 6,718 |
| Start/Stop codon-disrupting | 345 | 40 | 54 | 45 | 484 | 452 |
| **Total** | **5,987** | **279** | **4,766** | **6,999** | **18,031** | **13,031** |
| **Total number of genes affected by LOF variants\*** |  |  |  |  |  | **10,662** |

\* Some of the genes were affected with more than one LOF mutation, therefore the total number of genes is lower than the sum of the all genes.

To assess the quality of this catalogue of mutations, we first inspected it for genes already known (i.e. functionally validated) to harbor an LOF mutation. This is indeed the case, all known genes in the literature were found within the catalogue (**Supplementary data 6**). Then we investigated and confirmed the phenotypic impact of some of these LOF mutations in GmHapMap accessions (**Figure 4**). A frameshift mutation (frequency=0.003) in the microsomal omega-3 fatty acid desaturase (*FAD3A*), a key gene for linolenic acid synthesis in soybean seeds (Reinprecht & Pauls 2016), was found in three accessions. Near-infrared spectroscopy (NIRS) analysis of four soybean lines (two with and without this LOF mutation) showed a significant (*P* < 0.01) decrease in linolenic acid content in the mutant lines (4%) compared to the wild type (10%) (**Figure 4A**). A mutation (f=0.005) in *Glyma.04G050200*, the gene underlying the J locus controlling the Long Juvenile trait (Lu et al. 2017), resulted in a significant difference (*P* < 0.01) in grain weight per plant (8g in the mutant compared to 25g in the wild type) (**Figure 4B**). The introduction of a premature stop codon (f=0.02) due to a SNV in *GmGIa/E2* (Watanabe et al. 2012) significantly (*P* < 0.01) reduced the number of days from emergence to the appearance of the first open flower (DAE) (from 125 in wild-type lines to 95 in the mutant) (**Figure 4C**). Finally, a SNV (f=0.009) resulted in the disruption of splicing in the gene coding for the 3-ketoacyl-ACP synthase II (KASII) enzyme, a key gene in the oil biosynthesis pathway (Goettel et al. 2016). NIRS analysis of palmitic acid levels showed a significant (*P* < 0.05) decrease in the mutant lines (9%) compared to the wild type (12%) (**Figure 4D**). The development of a catalogue of LOF mutations represents a valuable resource for functional genomics.

**Figure 4.**
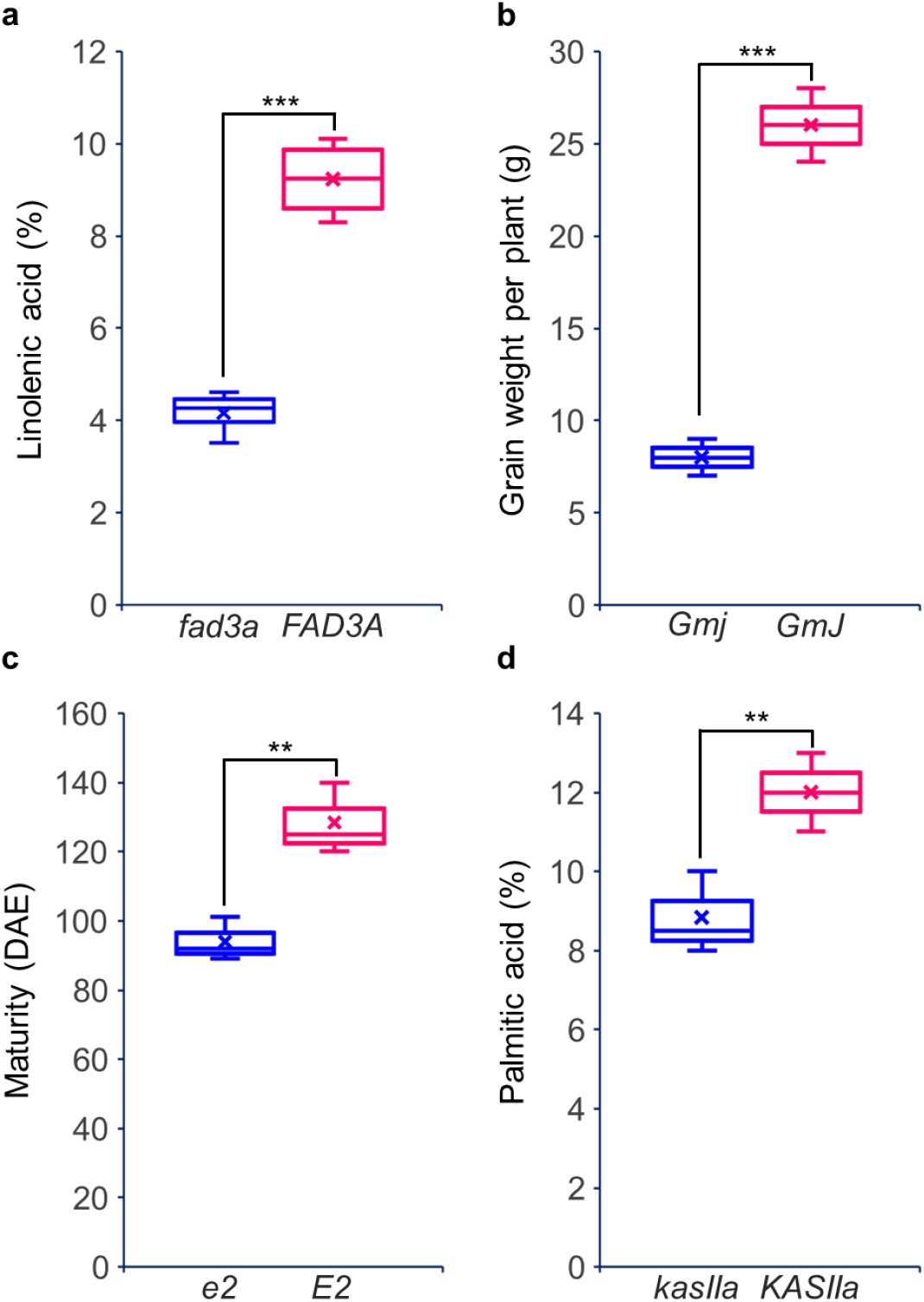
Phenotypic variation observed between accessions with (blue) and without (red) a predicted LOF mutation in four different genes. (**a**) *FAD3A*, a key gene for linolenic acid synthesis; (**b**) *GmJ*, a key gene of Long Juvenile trait; (**c**) *GmGIa*, a key gene controlling maturity; (**d**), *KASIIa*, a key gene in the oil biosynthesis pathway.

## Discussion

Using whole-genome sequencing data from a large collection of 1,007 soybean accessions, we developed the first haplotype map of soybean (GmHapMap), a valuable resource for soybean genetic studies and breeding. A first challenge was to create a uniform and accurate catalogue of nucleotide variation using a common version of the reference genome and a single bioinformatics pipeline (Lek et al. 2016). The GmHapMap produced here is not only uniform but also it achieved higher levels of genotype accuracy (>98%) compared to previous studies (92-97%) (Hwang et al. 2015). To create a representative haplotype map, a good level of saturation of both variants and haplotypes is required. Close to 15M sequence variants (SNVs, MNVs, and Indels) were called that captured nearly all polymorphisms with MAF > 1% in the USDA *G. max* germplasm collection (Song et al. 2013). The number of sequence variants did not increase significantly beyond the first 600 accessions, suggesting that a collection of this size has succeeded in capturing a sizeable fraction of worldwide nucleotide variation within cultivated soybean. Similarly, the number of unique haplotypes (4.3M tag SNPs) also plateaued relatively early within this collection of soybean germplasm. Together, these data suggest that the 15M variants captured in GmHapMap are both highly accurate and comprehensive of the genetic diversity within cultivated soybean at a worldwide level.

GmHapMap brings more resolution to the within-species diversity of *G. max*. A lower level of genomewide genetic diversity was observed here in soybean (mean θ_π_ = 1.36 × 10^−3^) compared to other major crops such as rice (θ_π_ = 2.29 × 10^−3^) (Caicedo et al. 2007) and corn (θ_π_ = 6.6 × 10^−3^) (Gore et al. 2009). It is presumed that several genetic bottlenecks, as well as strong selection pressure have reduced genetic diversity in soybean (Hyten et al. 2006). In addition, modern soybean breeding is founded on a very limited number of the founder accessions (Hymowitz et al. 1983). We also noticed an average Nonsyn/Syn ratio of 1.49, which is higher than that reported in other plants (sorghum (1.0), rice (1.2) and Arabidopsis (0.83) (Clark et al. 2007; McNally et al. 2009; Wang et al. 2015)). The greater accumulation of deleterious mutations in the soybean genome could be attributed to (1) a reduced effective population size (Makino et al. 2018); (2) a higher level of LD and the resulting ‘hitchhiking’ effect (Stephan et al. 2008); and (3) the domestication-associated Hill-Robertson effect (Lu et al. 2006).

The GmHapMap was used as a reference panel and more than 21K accessions that had been previously genotyped using common approaches (SNP array and/or GBS) and obtained an imputation accuracy of 92-96% for common variants and ~80% for rare variants. The accuracy levels, obtained here, are comparable to the 98% reported by Bukowski et al. (2018) in maize (Bukowski et al. 2018). The success of untyped-genotype imputation depends critically on how well a reference panel has captured the relevant haplotype diversity, as well as the marker density of the experimental dataset (Browning & Browning 2016). Here we document that GmHapMap provides an extensive capture of SNP and haplotype diversity within cultivated soybeans worldwide. It is likely that the lower imputation accuracy observed for the SNP array dataset can be attributed to the relatively low marker density of this dataset.

Enhanced datasets resulting from large-scale imputation can improve the efficacy of GWAS analysis (Hao et al. 2009; Marchini & Howie 2010). To illustrate the benefits of the GmHapMap resource for GWAS, we performed an association analysis on soybean seed oil content using imputed SNPs. A strong association with an imputed SNP residing in the *NPC1* gene was detected and its orthologue in Arabidopsis is known to contribute to seed oil content (Feldman et al. 2015). Several studies in human (Li et al. 2009), cattle (Santana et al. 2014), pig (Yan et al. 2017), maize (Yang et al. 2014) and rice (Wang et al. 2018) have demonstrated the capacity of imputation to improve the power of GWAS analysis. In the coming years, we expect that soybean researchers will deploy GmHapMap for imputation and more precise dissection of the genetic basis of complex traits in soybean.

This is the first time that a comprehensive description of GCHs, for the complete set of genes (55,589), has been achieved for a species. This catalogue of GCHs was obtained using HaplotypeMiner (Tardivel et al. unpublished). Tardivel et al. reported that HaplotypeMiner allowed the identification of SNP haplotypes for which 97.3% of lines sharing a same haplotype were correctly identified as having the same allele (Tardivel et al. unpublished). It has been well documented that haplotypes are more informative than single biallelic SNPs (Stephens et al. 2001). Knowledge of the GCHs (and possibly alleles) can greatly facilitate the establishment of a functional link between the various alleles of a gene and the associated phenotype. Haplotype-phenotype association revealed the functional alleles of several genes in wheat (Jiang et al. 2015), maize (Yang et al. 2013), rice (Si et al. 2016) and soybean (Langewisch et al. 2014). Knowledge of the alleles present at one or many genes can be tremendously important to breeders. Epistatic interactions between specific alleles as well as the effects of alleles at neighboring loci (carried along via linkage drag) can be very important when considering which combinations of alleles will be most desirable to achieve a given phenotype.

A final aspect of this work is that the identification of LOF mutations in soybean protein-coding genes. GmHapMap includes a set of nearly 11K knocked-out genes. We recognized that this catalogue of knocked-out genes is highly advantageous for soybean functional genomics for investigation of gene function, and application as genetic makers in soybean breeding programs.

The next challenge will be to link genetic variation, GCHs, and LOFs derived from GmHapMap with agronomic traits. This will need an extensive effort to measure phenotypes under multiple field and laboratory conditions. We believe that GmHapMap will lead and accelerate the soybean breeding efforts and future sustainable agriculture.

## Methods

### GmHapMap sequencing data

Two collections of soybeans were used: a first set of 727 accessions for which whole-genome sequencing had been previously released (Zhou et al. 2015; Maldonado dos Santos et al. 2016; Valliyodan et al.2016; Fang et al. 2017; Song et al. 2017; Torkamaneh et al. 2017) and a second set of 280 accession which were sequenced in this study. These were chosen to provide a more balanced representation of various soybean growing areas in the world. Seeds were planted in individual two-inch pots containing a single Jiffy peat pellet (Gérard Bourbeau & fils inc. Quebec, Canada). First trifoliate leaves from 12-day-old plants were harvested and immediately frozen in liquid nitrogen. Frozen leaf tissue was ground using a Qiagen TissueLyser. DNA was extracted from approximately 100 mg of ground tissue using the Qiagen Plant DNeasy Mini Kit according to the manufacturer’s protocol. DNA was quantified on a NanoDrop spectrophotometer. Illumina Paired-End libraries were constructed for 280 accessions using the KAPA Hyper Prep Kit (Kapa Biosystems, Wilmington, Massachusetts, USA) following the manufacturer’s instructions (KR0961 – v5.16). Samples were sequenced on an Illumina HiSeq X10 platform at the McGill University-Génome Québec Innovation Center in Montreal, QC, Canada.

### Nucleotide variants identification

Sequencing reads from all 1,007 accessions were processed using the same analytical bioinformatics pipeline (Fast-WGS) (Torkamaneh et al. 2017) to create a uniform catalogue of genetic variants. In brief, the 100-150-bp paired-end reads were mapped against the *G. max* reference genome [Gmax_275 (Wm82.a2)] (Schmutz et al. 2010). Then we removed variants if: 1) they had more than two alleles, 2) an allele was not supported by reads on both strands, 3) the overall quality (QUAL) score was <32, 4) the mapping quality (MQ) score was <30, 5) read depth (minNR) was <2 and 6) the minor allele frequency (MinMAF) was <0.0009.

### Determining the accuracy of nucleotide variants

The SoySNP50K iSelect BeadChip has been used to genotype the entire USDA soybean germplasm collection (Song et al. 2013). The complete dataset for 20,087 *G. max* and *G. soja* accessions genotyped with 42,508 SNPs was downloaded from Soybase (Grant et al. 2010). Of these accessions, we randomly selected 50 accessions which were in common with the GmHapMap collection. For these 50 accessions, we extracted their genotype calls at all SNP loci for which data were available. This large set of SoySNP50K genotype calls (2,125,400 genotypes or data points) was directly compared with the WGS-derived SNP calls (obtained using the Fast-WGS pipeline) to assess genotype accuracy.

### Determining the effects of nucleotide variants

The functional impact of nucleotide variants was performed using the soybean genome using SnpEff and SnpSift (Cingolani et al. 2012). Based on the genome annotation, nucleotide variants were categorized on the basis of their location (exonic, intronic, splice sites, UTR (3 & 5 prime), upstream and downstream regions (within 5kb of a gene), and intergenic) and their predicted functional impact (missense, nonsense, and silent). To determine LOF mutation, a database was built using 55K soybean protein-coding genes (Gmax_275_Wm82.a2.v1.gene.gff3, from Phytozome on Jan. 2016) for SnpEff. The LOF variants were extracted from the SnpEff-annotated VCF file using *grep* command lines. Variants were mapped on to transcripts annotated as “protein_coding” and containing an annotated “START” codon, and then classified as synonymous, missense, nonsense (stop codon-introducing, start/stop codon-disrupting or splice site-disrupting (canonical splice sites)). In this work, we excluded transcripts labelled as NMD (predicted to be subject to nonsense-mediated mRNA decay). We also applied another filtering step, based on annotation, to identify high-confidence knocked-out genes. The genes with LOF mutations were removed if (i) the ‘REF’ field in the input VCF file did not match the reference genome, (ii) they had an incomplete transcript, or (iii) they did not have a proper START codon.

### Population structure and genetic diversity

#### Structure

Population structure was estimated using a variational Bayesian inference implemented in fastSTRUCTURE (Raj et al. 2014). Five runs were performed for each number of populations (K) set from 1 to 15 using genomewide SNP data. The most likely K value was determined by the log probability of the data (LnP(D)) and delta K, based on the rate of change in LnP(D) between successive K values. Similar analyses were performed separately for all 20 chromosomes of soybean.

#### Genetic relationship

The evolutionary history was inferred using the Neighbor-Joining method (Saitou & Nei 1987) (rooted and unrooted) with the 12M genomewide SNPs identified in this study. The taxa were clustered together using bootstrap test (1,000 replicates) (Felsenstein 1985). The tree was drawn to scale, with branch lengths (next to the branches) in the same units as those of the evolutionary distances used to infer the phylogenetic tree. The evolutionary distances were computed using the Maximum Composite Likelihood method (Tamura et al. 2004) and the units correspond to the number of base substitutions per site. Evolutionary analyses were conducted in MEGA7 (Kumar et al. 2016).

#### Genetic diversity

We measured the nucleotide diversity (*π*) in sliding windows of 1000 bp across the genome using VCFtools (Danecek et al. 2011). The average pairwise divergence within a subpopulation (θ_π_) was estimated for the whole genome among different subpopulations. Sliding windows of different sizes (1 kb, 7kb, 10 kb and 100 kb) that had a 90% overlap between adjacent windows were used to estimate θ_π_ for both whole genome and each chromosome. To display the pattern in each chromosome, a window of 100 kb was used.

### Linkage disequilibrium and tag SNP identification

Genomewide pairwise linkage disequilibrium (LD) analysis (r^2^ and D’) was performed using all nucleotide variants from the GmHapMap dataset. The average r^2^ value was calculated for each length of distance (<1000 bp), and LD decay calculated using PopLDdecay (Zhang et al. 2018). For tag SNP selection, we used PLINK (Purcell et al. 2007) to calculate LD between each pair of SNPs within a sliding window of 50 SNPs and we removed all but one SNP that were in perfect LD (LD = 1); the remaining SNPs were deemed tag SNPs.

### Phasing, identification of identity by descent (IBD) and imputation

The IBD analysis was conducted using BEAGLE v4.1 (Browning & Browning 2016). In brief, to identify IBD segments, the genotypic dataset was phased using BEAGLE with 50 iterations for each chromosome. The output of these calculations was a series of “putative” IBD segments shared between pairs of individuals. Each segment comes with the following information attached: IDs for the pair of individuals, start and end position of the IBD segment, and probability score (LOD score). We filtered these segments using LOD score and the length of IBD.

### Imputation of untyped variants

We used two reference panels for untyped-variant imputation. The ‘REF-I’ panel includes 1,006 accessions from GmHapMap with the entire SNP dataset, while the ‘REF-II’ panel includes 1,006 accessions and only 1.9M tag SNPs from genic regions (tag SNPs in genic regions or within 2kb of a gene). These two reference panels were created for all 20 chromosomes of soybean and were phased using BEAGLE v4.1 (Browning & Browning 2016) with 100 iterations.

As initial lower density datasets, we used three collections of soybean accessions genotyped with commonly used genotyping tools. A first set of 20,087 accessions (the entire USDA Soybean Germplasm Collection) had been characterized using the SoySNP50K iSelect Bead Chip (Song et al. 2013) to yield a set of 43K polymorphic markers. A second set comprised 1,531 accessions which had been subjected to genotyping-by-sequencing (GBS; *Ape*KI protocol) (Sonah et al. 2013) and in which SNPs had been called using the Fast-GBS pipeline (Torkamaneh et al. 2017). Finally, a third set of 1,531 accessions (GBS set) with a combined SNP catalogue derived from GBS and SoySNP50K (**Supplementary Note**).

Phasing and imputation were performed using BEAGLE v4.1 (Browning & Browning 2016) for each chromosome with the following parameters: (i) nthreads = 10 (number of threads); (ii) window = 100,000 (number of markers in a sliding window); (iii) overlap = 50,000 (number of overlapping markers between adjacent windows); (iv) niterations = 100 (number of phasing iterations) and (v) err = 0.00001 (the allele miscall rate).

### Determining the imputation accuracy

The WGS SNP data from 1,006 of the 1,007 resequenced accessions were used as a reference panel to impute untyped variants. The remaining line was kept out of the reference panel to determine how accurately data at untyped loci (present in the GmHapMap data but absent from the low-density genotype catalogue) could be imputed in this accession. We performed three such permutations where a single accession was kept aside to estimate imputation accuracy. For these lines purposely excluded from the reference panel, we compared the imputed genotypes against the genotypes called at these same loci following WGS.

### Genomewide association analysis

Sonah et al. (2013) described a set of QTLs using GWA analysis on a subset of 139 soybean accessions. These accessions were genotyped via GBS. We imputed untyped variants on this low-density genotype dataset from GmHapMap in 1Mb of chromosome 14, encompassing a QTL for seed oil content. GWA analysis was conducted using GAPIT R package (Lipka et al. 2012) using a MLMM model (Segura et al. 2012). A candidate gene was identified using SoyBase database (Grant et al. 2010) and The Arabidopsis Information Resource (TAIR) [https://www.arabidopsis.org/servlets/TairObject?type=gene&name=AT4G38350.1]

### Identification of gene-centric haplotypes

The identification of GCHs was performed using the HaplotypeMiner R package (https://github.com/malemay/HaplotypeMiner) with the entire SNP dataset on 55,381 protein-coding genes in the soybean genome. In brief, the following parameters were used: (i) R2_measure = “r2s” (the estimation of linkage disequilibrium between markers was measured based on corrected *r^2^_vs_* which takes into account information related to genetic relatedness and population structure); (ii) cluster_R2 = “r2s” (LD measure to use in the clustering step); (iii) max_missing_threshold = 0.05 (the maximum proportion of missing genotypes allowed for a marker); (iv) max_het_threshold = 0.01 (the maximum proportion of heterozygous genotypes allowed for a marker); (v) min_allele_count = 4 (the minimum number of times the minor allele has to be seen for a marker to be retained); (vi) cluster_threshold = 0.9 (the minimum LD beyond which markers were clustered); (vii) max_flanking_pair_distance = 10000 (the maximum distance (in bp) that can separate two markers in LD at the final selection step: (viii) max_marker_to_gene_distance = 6000 (the maximum distance (in bp) from a marker to the center of the gene of interest); (ix) marker_independence_threshold = 0.8 (the minimum LD for two markers to be considered in LD at the final selection step).

### Identification of duplicated genes

We detected putative duplicated genes, presumably derived from WGD or gene duplication, using protein homology analysis integrated in the Phytozome (Goodstein et al. 2012) and SoyBase (Grant et al. 2010) databases. Protein homologs were identified using dual-affine Smith-Waterman alignments between the predicted translation product of the selected transcript (aka query gene) and all other predicted proteins in the soybean genome. We identified duplicated genes with 90% identity (ID≥90), 90% coverage (CV≥90), and 5% size difference (SD≤5) threshold.

### Code availability

The bioinformatics codes and scripts applied in this study for variant calling, population structure analysis, genetic diversity, tag SNP selection, imputation, GWAS, annotation and GCHs detection are publicly available at https://figshare.com/account/home#/projects/56921.

### Data availability

The datasets produced in this study (GmHapMap nucleotide variants (complete dataset), reference panels (REF-I and REF-II), annotated GmHapMap nucleotide variants, GCHs for all 55K genes and LOF variants and genotypes) are publicly available at https://figshare.com/account/home#/projects/56921

## Acknowledgments

This work was supported by the SoyaGen grant (www.soyagen.ca) awarded to F. Belzile and funded by Génome Québec, Genome Canada, the government of Canada, the Ministère de l’Économie, Science et Innovation du Québec, Semences Prograin Inc., Syngenta Canada Inc., Sevita Genetics, Coop Fédérée, Grain Farmers of Ontario, Saskatchewan Pulse Growers, Manitoba Pulse & Soybean Growers, the Canadian Field Crop Research Alliance and Producteurs de grains du Québec. The work conducted by the US DOE Joint Genome Institute is supported by the Office of Science of the US DOE under Contract DE-AC02-05CH11231. We thank Dr. Brain Boyle to help for DNA library construction. We thank the Joint Genome Institute and collaborators for pre-publication access to the *Glycine max* W82 V2 genome sequence. We also thank Dr. Gary Stacy and Dr. Shakhawat Hossain for pre-publication access to the soybean transcriptome atlas.

## Author contributions

DT and FB conceived the project. DT and JL contributed to programming and analysis of genomic data. DT carried out genetic diversity analysis, imputation, GCHs and identification of LOFs. BV and HN provided sequence data for American accessions. RA provided data for Brazilian accessions. DT, FB, EC, IR and LO provided data for Canadian accessions and also carried out NIRS analysis. AS and JS carried out the identification of gene duplication. DT and FB contributed to writing the manuscript.

